# Short linear motifs - Underexplored players driving *Toxoplasma gondii* infection

**DOI:** 10.64898/2026.01.12.699020

**Authors:** Jesús Alvarado Valverde, Karine Lapouge, Arne Boergel, Kim Remans, Katja Luck, Toby J. Gibson

## Abstract

Pathogens infect hosts by interacting with host proteins and exploiting their functions to their advantage. Short linear motifs, small functional regions within intrinsically disordered protein regions, are common mediators of host-pathogen protein interactions. While motifs have been more extensively studied in viruses and bacteria, the extent to which eukaryotic unicellular parasites use motifs during infection remains unexplored. *Toxoplasma gondii* is a widespread intracellular Apicomplexan parasite capable of infecting all warm-blooded animals and invading any of their nucleated cells. *Toxoplasma’s* secreted proteins are key in interacting with host proteins during infection, making them potential sources for motifs. To highlight the role of motifs in *Toxoplasma gondii* infection, we curated 21 known motif instances in Toxoplasma proteins from the scientific literature. To identify more motifs in Toxoplasma secreted proteins, we developed a computational pipeline that annotates putative motif matches with structural and functional features. Through this approach, we identified a set of 24,291 motif matches in 295 secreted proteins. We highlight strategies for further prioritisation of likely functional motif matches by focusing on integrin motifs, degrons and TRAF6-binding motifs. We subjected four predicted TRAF6-binding motifs to experimental validation, supporting the predicted motifs in the Toxoplasma proteins RON10 and GRA15. Our motif predictions provide a valuable resource for generating hypotheses and designing experiments to study infection mechanisms. The characterisation of motifs in *Toxoplasma* will be key to understanding the molecular principles underlying its broad host range and more comprehensive Apicomplexan infection strategies.

**Importance:** Toxoplasma gondii is a widely distributed intracellular parasite that achieves a successful infection by interacting with different host cell proteins. Short linear motifs are small functional modules found in unstructured protein regions and recognised by folded protein domains. Given that unstructured protein regions are a common feature of Toxoplasma’s proteins, we hypothesise that motifs play important roles during its infection cycle. Here, we highlight the role of motifs during the Toxoplasma host cell invasion cycle through a curated set of motif examples. Through a computational pipeline, we predict thousands of motifs in secreted proteins, outline strategies for working with these predictions and finally experimentally test proteins containing a motif involved in the innate immune response, successfully showing the binding of two motifs. Our work provides a resource for further motif testing in Toxoplasma proteins, aiming at understanding the molecular mechanisms of its infection strategies and its broad host range.

## Introduction

Intracellular pathogens, such as viruses, bacteria or eukaryotic parasites, interact with host cell components throughout their infection cycles. These pathogens can attach to the host membrane, and when internalised, they can interact with the host cytoskeleton and organelles, as well as with its export and translation machinery (1–3). By interfering with host biological processes, pathogens modulate the host immune response, replicate and disperse. Pathogen proteins play a central role in these interactions. A common mode of protein binding that mediates host-pathogen protein-protein interactions (PPIs) consists of a folded domain, e.g. in host proteins, and short linear motifs (hereafter referred to as motifs) in pathogen proteins (2–4). Motifs are usually between 3 and 10 amino acids long and occur in intrinsically disordered regions of proteins (IDRs)(5,6). Motifs mediate protein interactions that primarily participate in regulatory and cell signalling processes. Motifs define sites for post-translational modifications, subcellular protein localisation, as well as substrate recognition signals for enzymes such as E3 ligases, proteases, phosphatases, etc (5). Motifs are unstructured in their unbound state and often adopt an alpha-helix or beta-strand upon binding to partner proteins (6). Because only a few mutations are needed to create a motif, they can evolve in much shorter evolutionary timescales compared to folded protein regions (7). Thus, pathogens have evolved to exploit these host motifs to infect their hosts, probably because motifs are short and easily emerge in proteins (7), but also because of their many functional roles in cell signalling. Many instances of host motif mimicry have been characterised for viruses and bacteria (2,3,8–11). Although there have been some studies focusing on intracellular eukaryotic parasites (12,13), the extent to which parasites use motifs is less understood. *Toxoplasma gondii,* a member of the Apicomplexa phylum of unicellular parasites, has one of the proteomes with higher levels of IDRs among Eukaryotes (14,15), making it a good candidate to discover and characterise the role of motifs in its proteins during parasitic infection.

*Toxoplasma* is a successful eukaryotic unicellular parasite, as it can infect any warm-blooded animal and proliferate within any of its nucleated cells. It is estimated that a third of the human population has been infected with Toxoplasma (16). Even though most of the infections in humans are asymptomatic, immunocompromised individuals and susceptible populations can develop different forms of Toxoplasmosis, from influenza-like symptoms to severe phenotypes such as encephalitis, making *Toxoplasma* a global health burden (16,17). *Toxoplasma’s* definitive hosts are cats and all members of the feline family, where it reproduces sexually in their gut to then disseminate in the environment. In all other hosts, such as all mammals and birds, *Toxoplasma* reproduces asexually in different cell types to multiply its numbers and disseminate along the food chain (18). Having multiple hosts means that *Toxoplasma* not only has the molecular tools to infect different host cells but also to deal with diverse cell programmes and immune responses, to thrive in each of its hosts. Motifs in Toxoplasma proteins might play important roles in mediating the adaptation to the many different host environments this parasite encounters.

Once consumed by its host, *T. gondii* cell invasion cycles depend on the sequential secretion of proteins contained in its specialised infection organelles: the micronemes, rhoptries and dense granules (Fig. 1A). Microneme proteins, or MICs, are targeted to the parasite cell surface and are involved in host recognition and attachment, while rhoptry proteins, termed RONs and ROPs, are secreted into the host cytoplasm and are involved in host invasion and niche establishment (Fig. 1B). During the invasion process, a parasitophorous vacuole (PV) is created from the invagination of the host cell membrane (19). From within this space, *Toxoplasma* creates a suitable environment and continues to secrete proteins from membraneless organelles called dense granules. These proteins, mostly known as GRAs, are secreted either into the PV lumen, inserted into its membrane (PVM) or into the host cytoplasm and even into the host nucleus, helping the parasite in modulating the host cell’s regulatory processes to proliferate (20). All these secreted proteins interact with host proteins throughout the infection cycle, including host cell receptors, cytoskeletal proteins and transcription factors (21,22). These host-parasite protein interactions are the subject of continuous study, serve as a reference to study other Apicomplexans, and are promising targets for the development of therapies. Knowledge about motifs in *Toxoplasma* secreted proteins would increase our mechanistic understanding of Toxoplasma infection strategies.

**Figure 1.**
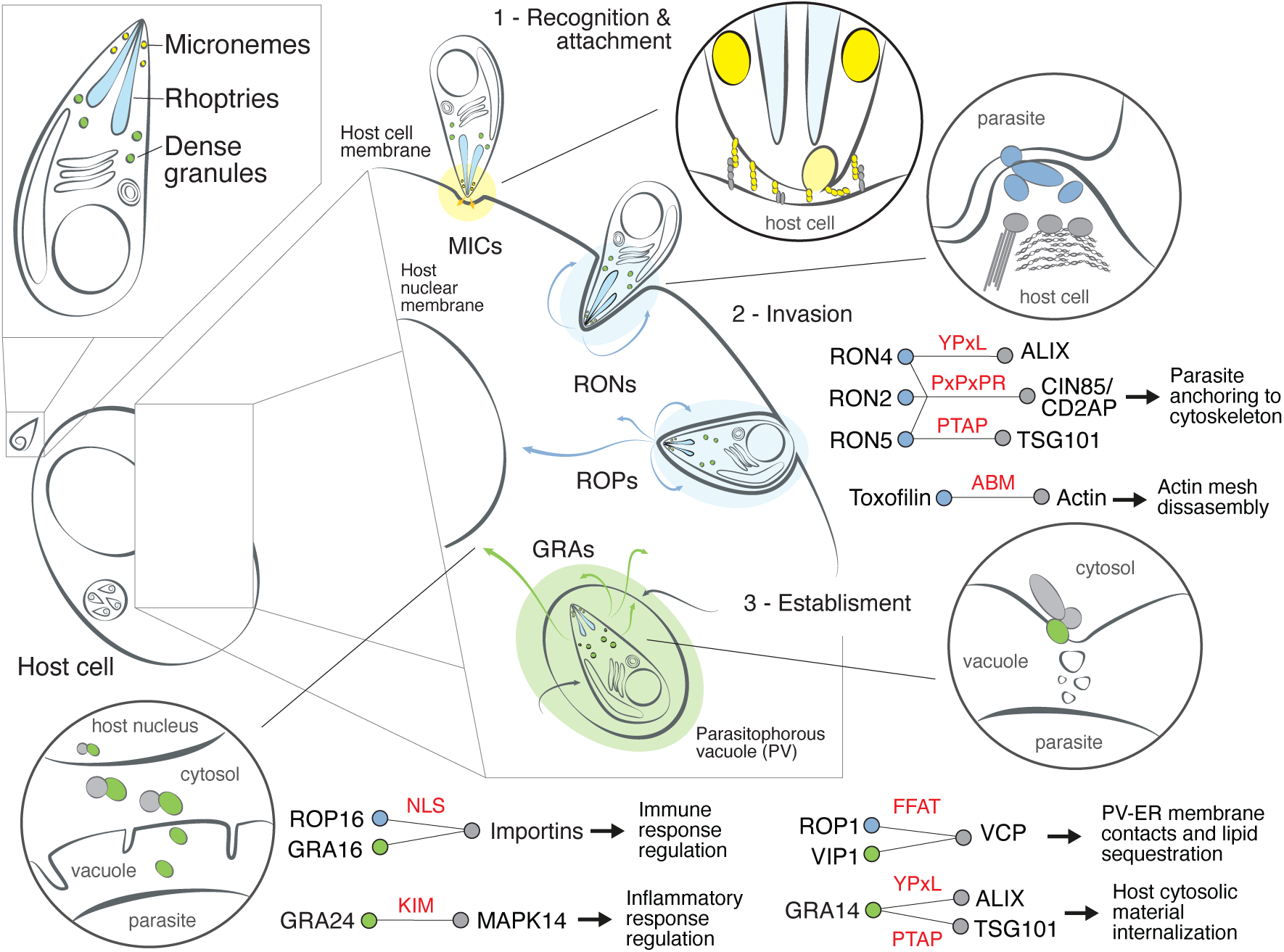
Motif use and hijacking during the *Toxoplasma* infection process. *Toxoplasma gondii* secretory organelles and infection cycle illustrating different host-pathogen interaction contexts, known motif instances and their infection outcomes. Motif names are highlighted in red, proteins from micronemes are highlighted in yellow, those from rhoptries in blue, and those from dense granules in green.

To identify motifs in *Toxoplasma gondii* proteins, we collected motif instances in *T. gondii* from the literature, predicted occurrences of known motif types in its secreted proteins through the development of an integrative discovery pipeline, and tested the binding of predicted TRAF6 binding motifs *in vitro,* which might have relevance in the regulation of innate immune response. The available motif predictions along with annotations will motivate further functional studies in *Toxoplasma* and other Apicomplexan parasites to understand mechanisms of infection and provide further examples of motif repurposing among intracellular pathogens.

## Results

### Curation of known motifs in *Toxoplasma gondii* secreted proteins

We collected 21 experimentally validated motif instances from 11 Toxoplasma proteins from the scientific literature that serve as a reference of the current motif knowledge in *Toxoplasma gondii* (Table 1, Fig. 1). The Eukaryotic Linear Motif (ELM) resource is a reference point for curated eukaryotic motifs from the scientific literature. Based on motif sequence patterns, structures, binder information, and functional annotations, curators define motif types and group motif types into broader motif classes (23). Two of the Toxoplasma motif instances were previously annotated as substrate recognition sites in ELM (DOC). Eleven motif instances were novel with respect to ELM content and were added to known motif classes of scaffolding classes in ELM (LIG). The remaining eight novel motif instances are similar but were not able to be directly annotated to ELM. Those correspond to one scaffolding class (LIG) and to three subcellular localisation classes (TRG in ELM). We did not annotate motif instances in Toxoplasma secreted proteins reported as motifs of cleavage sites (CLV in ELM), degradation signals (DEG in ELM) or sites for post-translational modifications (MOD in ELM). As detailed below, the curated 21 motif instances function throughout parasite invasion and establishment (Fig. 1).

**Table 1.**
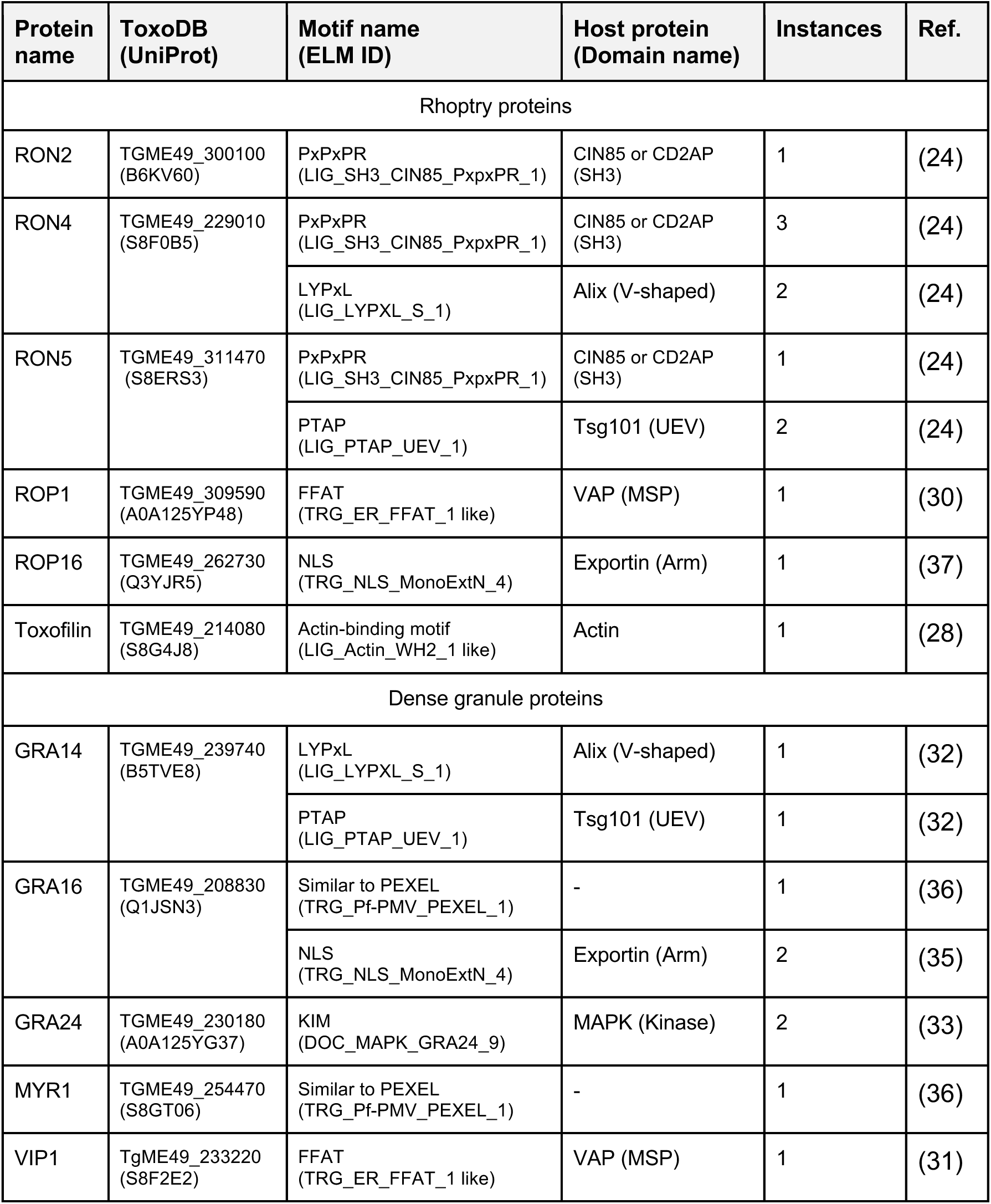
Experimentally validated linear motif instances. . Known *Toxoplasma* motif instances from the literature and interacting host protein domains. Protein IDs correspond to the ToxoDB and UniProt accessions, while motif IDs correspond to the ELM motif classes.

| Protein name | ToxoDB (UniProt) | Motif name (ELM ID) | Host protein (Domain name) | Instances | Ref. |
| --- | --- | --- | --- | --- | --- |
| Rhoptry proteins |  |  |  |  |  |
| RON2 | TGME49_300100 (B6KV60) | PxPxPR (LIG_SH3_CIN85_PxpxPR_1) | CIN85 or CD2AP (SH3) | 1 | (24) |
| RON4 | TGME49_229010 (S8F0B5) | PxPxPR (LIG_SH3_CIN85_PxpxPR_1) | CIN85 or CD2AP (SH3) | 3 | (24) |
|  |  | LYPxL (LIG_LYPXL_S_1) | Alix (V-shaped) | 2 | (24) |
| RON5 | TGME49_311470 (S8ERS3) | PxPxPR (LIG_SH3_CIN85_PxpxPR_1) | CIN85 or CD2AP (SH3) | 1 | (24) |
|  |  | PTAP (LIG_PTAP_UEV_1) | Tsg101 (UEV) | 2 | (24) |
| ROP1 | TGME49_309590 (A0A125YP48) | FFAT (TRG_ER_FFAT_1 like) | VAP (MSP) | 1 | (30) |
| ROP16 | TGME49_262730 (Q3YJR5) | NLS (TRG_NLS_MonoExtN_4) | Exportin (Arm) | 1 | (37) |
| Toxofilin | TGME49_214080 (S8G4J8) | Actin-binding motif (LIG_Actin_WH2_1 like) | Actin | 1 | (28) |
| Dense granule proteins |  |  |  |  |  |
| GRA14 | TGME49_239740 (B5TVE8) | LYPxL (LIG_LYPXL_S_1) | Alix (V-shaped) | 1 | (32) |
|  |  | PTAP (LIG_PTAP_UEV_1) | Tsg101 (UEV) | 1 | (32) |
| GRA16 | TGME49_208830 (Q1JSN3) | Similar to PEXEL (TRG_Pf-PMV_PEXEL_1) | - | 1 | (36) |
|  |  | NLS (TRG_NLS_MonoExtN_4) | Exportin (Arm) | 2 | (35) |
| GRA24 | TGME49_230180 (A0A125YG37) | KIM (DOC_MAPK_GRA24_9) | MAPK (Kinase) | 2 | (33) |
| MYR1 | TGME49_254470 (S8GT06) | Similar to PEXEL (TRG_Pf-PMV_PEXEL_1) | - | 1 | (36) |
| VIP1 | TgME49_233220 (S8F2E2) | FFAT (TRG_ER_FFAT_1 like) | VAP (MSP) | 1 | (31) |

Rhoptry proteins RON2, RON4 and RON5 each contain PxpxPR motifs. During host cell invasion, these motifs help the parasite connect its tight junction (TJ) complex to the host cytoskeleton (24) (Fig. 1). The PxpxPR motifs bind to the SH3 domain of host adaptor proteins CIN85 or CD2AP, which are involved in cytoskeletal rearrangements, vesicle trafficking and endocytosis (25,26). Additionally, RON4 and RON5 contain instances of the motif types LYPxL and PTAP. The LYPxL motif mediates the interaction of proteins with ALIX, while the PTAP motif binds to the UEV domain of class E vacuolar sorting protein TSG101. ALIX and TSG101 are both key components of the ESCRT membrane remodelling system, and during *Toxoplasma* cell invasion, they are also recruited to the TJ for an efficient cell entry (24,27). This process is also aided by parasite proteins such as Toxifilin, which binds to host actin through an actin-binding motif (ABM) that is similar in sequence pattern to the annotated WH2-binding motif in ELM (28). Toxofilin sequesters actin and promotes the disassembly of the host cortical actin meshwork, facilitating an efficient internalisation (29).

Once *Toxoplasma* is established within the host cell, ROP1 and dense granule proteins, VIP1 and GRA14 are secreted to the parasitophorous vacuole membrane (PVM) (Fig. 1). ROP1 and VIP1 both contain FFAT motifs (similar in sequence motif pattern to the FFAT motif type in ELM) that modulate the targeting of proteins to the endoplasmic reticulum via its interaction with Vesicle Associated Membrane Protein (VAMP) Associated Proteins (VAPs). As ROP1 and VIP1 are inserted in the PVM, their interaction with VAP proteins leads to the association of the PVM with the host ER membrane. Toxoplasma exploits this proximity to sequester the lipids necessary for its vacuole and plasma membrane biogenesis (30,31). GRA14 harbours both LYPxL and PTAP motifs in its C-terminal region. Since GRA14 is also inserted into the PVM with its C-terminus facing the host cytosol, the motifs can recruit ALIX and TSG101, which, in this case and differently from TJ proteins, help *Toxoplasma* sequester host cytosolic material through the budding of its PVM (32).

Further, during establishment, more parasite proteins are secreted into the host cytosol and may then be transported to the host nucleus. Within the host cytosol, secreted proteins interact with host proteins via motifs. The intrinsically disordered protein GRA24 contains two instances of a motif that binds to mitogen-activated protein kinase p38α (MAPK14). These motifs allow GRA24 to bind to a MAPK14 dimer, forming a stable complex that leads to the expression of proinflammatory factors, which in turn modulate parasite replication (33,34). Parasite protein export is achieved by the presence of cell compartment localisation motifs, which either use the parasite or host machinery to be secreted. In the first case, the correct export of dense granule protein MYR1 to the PV and of GRA16 to the cytosol is dependent on the cleavage of a short linear motif by a parasite aspartyl protease, in a similar way as the Plasmodium PEXEL export motif (35,36). In the case of host nuclear targeting, GRA16, as well as rhoptry protein kinase ROP16, are imported into the host nucleus via nuclear localisation signals (NLS). ROP16 activates signal transducer and activator of transcription (STAT) signalling pathways to inhibit host cell death and modulate the production of interleukins (37,38). Within the nucleus, GRA16 regulates genes involved in cell cycle progression (35). This set of curated motif instances provides a non-exhaustive reference and exemplifies their role during important stages of the Toxoplasma invasion cycle. Nevertheless, the small number of known motifs as well as their occurrence in a few proteins, highlight the need and potential to find and characterise many more motif instances to facilitate further mechanistic studies of Toxoplasma infection strategies.

### Prediction of motif instances in *Toxoplasma* secreted proteins

To identify novel motif instances that function in the Toxoplasma infection process via interaction with host proteins, we focused on *Toxoplasma gondii* proteins from the strain ME49 that are secreted into host cells. To determine the subcellular localisation of secreted proteins, we used the dataset from the hyperLOPIT experiment (39) and annotations from the ToxoDB, a database from the VEuPathDB resource, the reference database for the research of eukaryotic pathogens (protists and fungi), their hosts and invertebrate vectors (40). The secreted proteins were subsequently filtered for those with evidence of expression using mass spectrometry datasets from the ToxoDB. To predict motif instances in these 314 secreted proteins, we developed a computational pipeline based on sequence pattern searches as well as structural and functional annotations of predicted motifs (Fig. 2). To find putative motif instances, we used regular expressions defined for annotated motif classes in the ELM database, which capture the preferences for certain amino acids at certain motif positions. We scanned the selected proteins with 298 regular expressions corresponding to motif classes with interacting domains that could be present either in Toxoplasma or in the host using the taxonomic range annotations provided by the ELM resource. This motif search resulted in a total of 68,066 motif matches in 309 proteins. Since motifs occur primarily in IDRs and need to be accessible for binding to the partner proteins, we deployed structural annotations to further restrict to more likely functional motif matches. To this end, we evaluated the location of the motif match in IDRs using the software IUPRed3(41). In addition, in cases where a full-length structural model of a given protein was available from the EBI AlphaFold database (42), we evaluated motif accessibility through low pLDDT values and high relative solvent accessibility (RSA). In cases where a structural model was not available, we excluded motif matches that fell within annotated folded domain boundaries as annotated in UniProt (43) (Fig. 2). Using this framework, we obtained a set of 24,291 motif matches within 295 proteins from micronemes, rhoptries and dense granules (Fig. 2, Fig. 3A) (Table S1).

**Figure 2.**
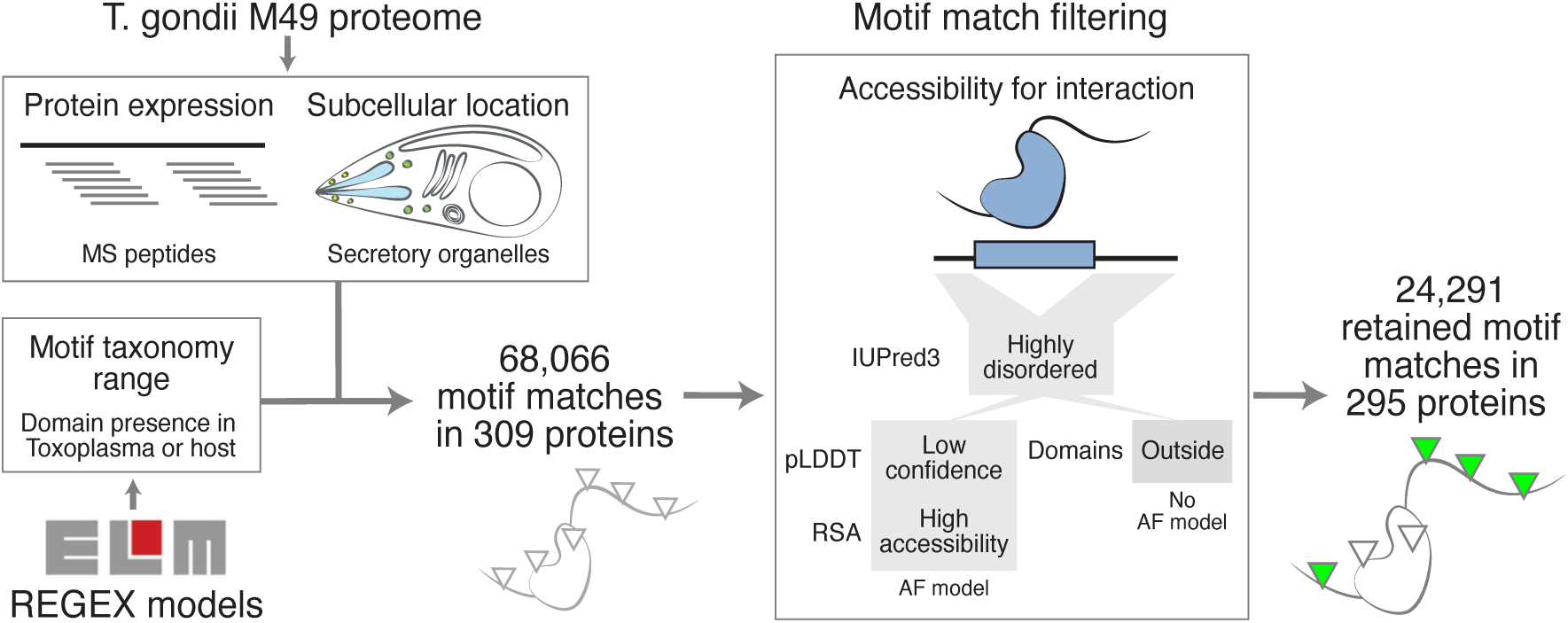
Motif prediction pipeline. Diagram outlining the protein and motif model selection for motif prediction, as well as motif accessibility filtering criteria.

**Figure 3.**
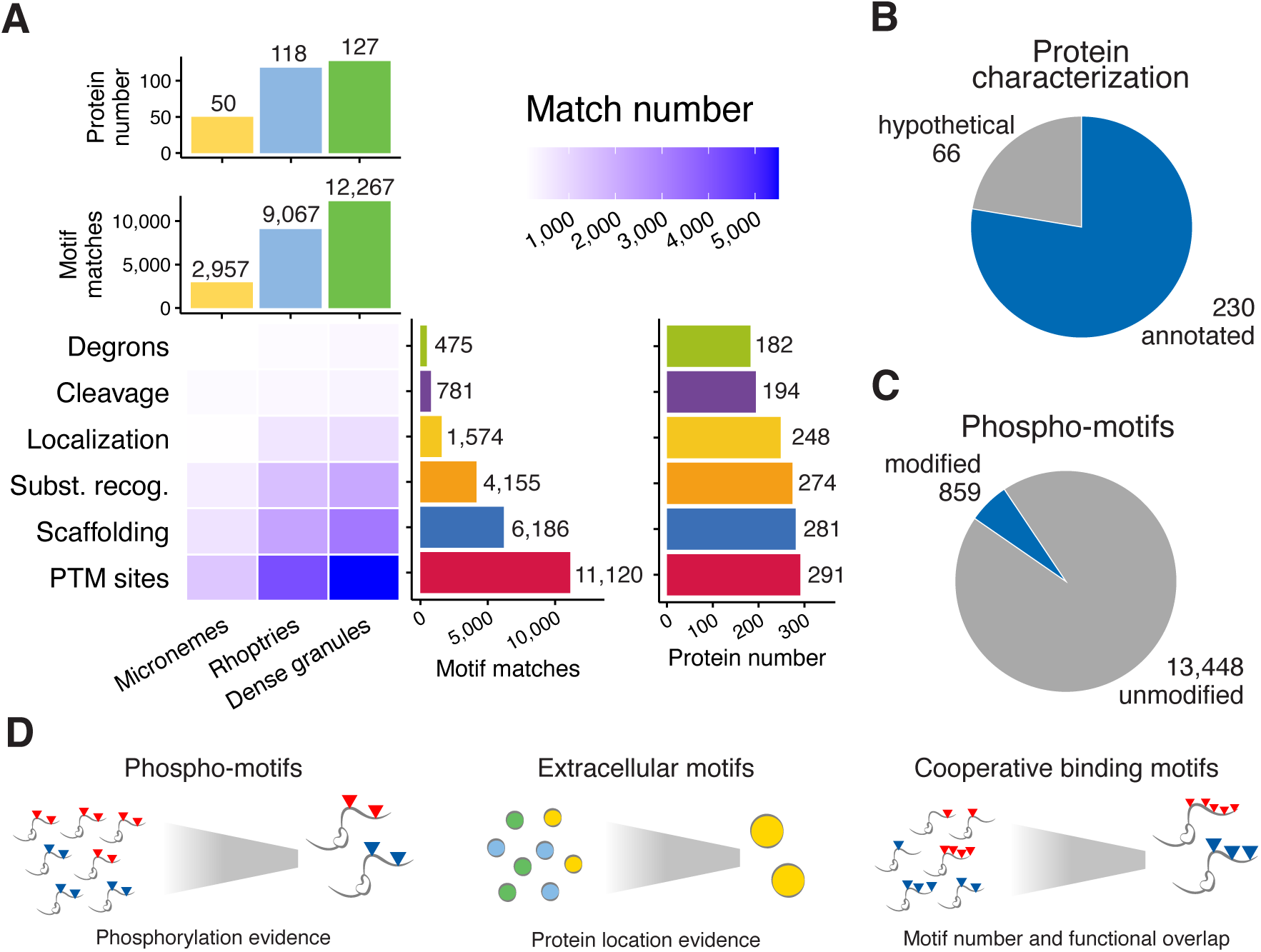
Matches in secreted proteins. **A**. Number of filtered motif matches from secreted proteins across different motif categories. **B**. Split of proteins with filtered matches according to their annotation. **C**. Split of filtered phospho-motifs according to whether they have phosphorylation evidence. **D**. Additional motif filtering strategies according to motif type functions and properties.

Our computational pipeline captured 16 of the 21 known motif instances within our retained motif matches. Two PTAP motifs in RON5 were missed because they have low protein IDR scores. The FFAT motifs in ROP1 and VIP1, and the actin-binding motif in Toxofilin were not captured because their sequences did not match any ELM regular expression. These last two examples highlight that current annotated motif types in the ELM resource do not fully capture the diversity of motifs in *Toxoplasma* proteins. Our curated motif instances could thus be used to reevaluate and extend ELM motif type annotations. When further analysing the curated motif instances here, we found a previously unreported PxpxPR motif in the N-terminal region of RON4 as well as two SxIP motifs. SxIP motifs modulate the binding to microtubule plus-end tracking proteins (+TIPs), which have an important role in chromosome segregation, cell signalling and organelle trafficking (36). SxIP motifs are already known to be exploited by the Apicomplexan parasite *Theileria* (13). RON8, which, together with RON2, RON4, and RON5, forms part of the TJ (22), also contained matches of the SxIP and LYPxL motifs. The presence of extra motif matches related to host cytoskeleton in TJ proteins indicates that the current model of motif-hijacking during this stage of host cell invasion is still incomplete and will benefit from our motif predictions. Furthermore, the identification of 4,822 strong motif matches in our dataset from 66 proteins annotated as “hypothetical” in ToxoDB and with expression evidence (Fig. 3B) represents a potential venue to explore, assign and understand their function.

On average, our pipeline obtained 82 retained motifs per secreted protein: 59 in micronemes, 77 in rhoptries and 97 in dense granules (Fig. 3A). While proteins, especially those with longer IDRs such as dense granule proteins (Fig. S1A), are expected to have multiple motif occurrences, the relatively high number of predicted motif instances still points to some of those being likely non-functional. Most of the motif matches come from motif types that denote sites for PTMs such as phosphorylation (Fig. 3A). This most likely results from very degenerate regular expressions available for motif types from the PTM class, such as …[ST]…[ST] defining sites phosphorylated by GSK kinase. In regular expressions, a “.” corresponds to any amino acid possible at this position and amino acids in brackets denote possible amino acids at this position in the motif. More than 2,000 motif matches alone were identified with this GSK kinase phosphorylation motif pattern. Phosphoproteomics data can be used to provide evidence for actual modification of these predicted motifs (45). Using available phosphosite datasets from ToxoDB, we found evidence of phosphorylation for ∼6% of predicted phosphorylation sites (Fig. 3C-D), assigning higher confidence to these predicted motifs (Table S1). We generally observe that for most motif classes, motif matches were identified only for a fraction of annotated motif types in ELM. For example, 38% of motif matches from the targeting class (TRG) correspond to diArg motifs, 35% of motif matches from the degradation class (DEG) correspond to SPOP degrons (see below), and 18% of scaffolding motifs bind SH3 domains (Fig. S1B).

To further select for more likely functional motif matches, one can take into account functional aspects of the different motif types and the cellular context of secreted proteins (Fig. 3D). For example, as microneme proteins will be present in extracellular environments, motifs known to interact with domains from extracellular proteins would be expected to have more likely functional relevance. In our dataset (Table S1), we found 401 motif matches from 12 motif extracellular types in 48 microneme proteins that could be relevant during the recognition and attachment stages. Integrin binding is a common strategy for pathogens to attach to the host cell membrane (46–48). The most common integrin-binding motif is the RGD motif, which can bind to eight of the different alpha and beta integrin heterodimers (49). Within our dataset, we found three motif matches of the RGD motif in three microneme proteins: MIC17C, Toxolysin 4 (TLN4) and chitinase-like protein CLP1. As some microneme proteins might be localised to the host cytosol, e.g. during host cell lysis and egress (50), we did not filter out matches from other cytosolic motif types in microneme proteins. Degradation signals, or degrons, are motifs recognised by E3 ubiquitin ligases and mark proteins for proteasomal degradation (51). Degrons might be relevant motif candidates for Toxoplasma infection, as other pathogens are known to modify protein levels via degrons to achieve outcomes such as the stabilisation of host components necessary for their survival (2) or modulating the activity of their proteins once inside the host (52). We found 475 degrons in 182 secreted proteins: 73 in micronemes, 157 in rhoptries and 245 in dense granule proteins. Speckle-type POZ protein (SPOP), an E3 ubiquitin ligase, recognises its substrates via an ST-rich degron (53). We found

155 SPOP degron matches in 80 secreted proteins. Since multivalency is a known mechanism by which SPOP substrates enhance their affinity to SPOP (53), one could prioritise the 34 secreted proteins containing two or more SPOP motif matches in our dataset. Rhoptry protein toxolysin 1 (TLN1), a metalloprotease with a role in rhoptry maturation (54), but potentially detrimental for host establishment once secreted into the host cytosol, has 10 identical SPOP degron matches, which could be used to modulate TLN1 levels once within the host cytoplasm. Overall, these examples illustrate how our dataset can be further explored to prioritise motif candidates for experimental characterisation.

### Experimental validation of predicted TRAF6-binding motifs

For the experimental validation of predicted motifs, we focused on motifs that bind the tumour necrosis factor (TNF) receptor-associated factor 6 (TRAF6) protein. TRAF6 binds to proteins containing a P.E..[DE(*ɸ*)] motif through its C-terminal MATH domain (*ɸ* denotes hydrophobic residues). TRAF6 plays important roles in immunity and development by contributing to the formation of various signalling protein complexes. During the innate immune response, TRAF6 is activated by interleukin 1 receptor (IL-1R), linking this receptor to the canonical activation of the transcription factor NF-κB (55). Through the addition of K63-linked polyubiquitin chains, TRAF6 primes proteins for complex assembly and not for proteasomal degradation. The TRAF6-binding motif is present in proteins such as IL-1R, Tumour necrosis factor protein CD40, and Mitochondrial antiviral-signalling protein (MAVS) (56). In the context of infection, herpesviruses have been shown to use the TRAF6 motif-containing protein Tio to stimulate NF-κB activation (57).

Among our predicted motif instances, we found 111 motif matches in 68 rhoptry and dense granule proteins. Given that the TRAF6-binding motif is known to appear multiple times within proteins (58), we hypothesised that the motif repetitions could interact with the TRAF6 trimer in a multivalent way. Following this criterion, we selected rhoptry neck protein RON6, which harbours 17 matches of the TRAF6-binding motif, most of them being identical repeats, and RON10, which harbours two different TRAF6-binding motif matches (Fig. 4A). RON10 is known to interact with RON9, but RON10 and RON9 do not have clear roles during infection (59). TRAF6 has been shown previously to interact with the dense granule proteins GRA7 and GRA15, which themselves play a role in modulating the innate immune response during *Toxoplasma* infection (60–63). However, the actual interaction interfaces of GRA7 and GRA15 with TRAF6 have not been described (22), making the TRAF6 motif matches in GRA7 and GRA15 interesting candidates for further experimental characterisation (Fig. 4A).

**Figure 4.**
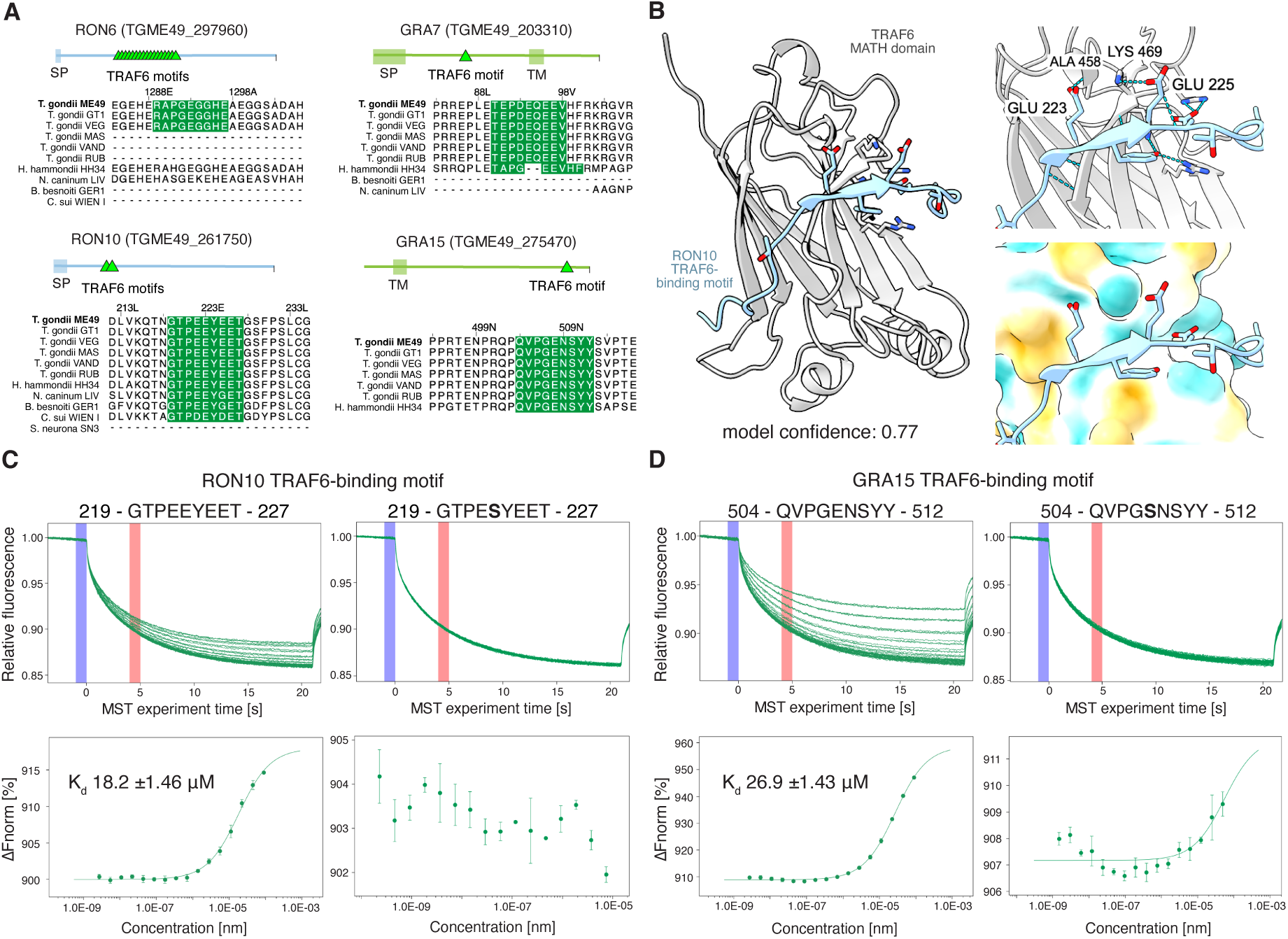
TRAF6 motif binding assays. **A.** Protein diagrams of RON6, RON10, GRA7 and GRA15 indicating the position of their TRAF6-binding motifs with accompanying multiple sequence alignment showing their presence in different Toxoplasma strains and Coccidian species. **B.** AlphaFold model of TRAF6 MATH domain with TRAF6-binding motif from RON10. Right panels display the potential hydrogen bond network at the interaction interface (top) and secondary structure adopted by the TRAF6-binding motif (bottom). **C.** RON10 TRAF6 motif testing. MST traces (top graph) and dose-response curves (bottom panel) of TRAF6 MATH domain binding to RON10 TRAF6 motif containing peptide (left) and the E>S mutant (right). In the MST traces, the cold region is set at 0 s (blue) and the hot detection region at 5 s (red). **D**. Same as in A, but for the GRA15 TRAF6 motif and the E>S motif mutant.

We proceeded to determine the dissociation constants of the TRAF6-binding motifs in RON6, RON10, GRA7 and GRA15 to the TRAF6 MATH domain using the microscale thermophoresis (MST) binding assay. To this end, we expressed and purified the human TRAF6 MATH domain and ordered peptides containing the sequence of the following motif matches. In the case of RON6, we selected the most frequent version of its motif repeats. For RON10, we selected the second motif match that contained multiple negative charges, which might contribute to binding (Fig. 4B). GRA7 and GRA15 only have one TRAF6-binding motif match each, which was selected for experimental testing. In addition, we ordered these wildtype peptide sequences with a Glutamate to Serine mutation at position 3 in the motifs supposed to weaken or abrogate binding to the MATH domain because the conserved glutamate residue forms a hydrogen bond with backbone atoms of residue A458 in TRAF6 (Fig. 4B) (<u>Table S2</u>). Our experimental results confirm that the peptides containing the motifs from RON10 and GRA15 can bind to the TRAF6 MATH domain, and these interactions were lost when testing the mutant peptide versions. The dissociation constants for the RON10 (Kd 18.2 ±1.46 μM) and GRA15 peptides (Kd 26.9 ±1.43 μM) are as expected, for functional motif interactions, in the lower micromolar range (Fig. 4C-D, <u>Data S1</u>, <u>Table S2</u>). These results represent the first evidence that RON10 can bind TRAF6, pointing to a role of RON10 in the regulation of the innate immune response.

## Discussion

It has been previously speculated that *Toxoplasma* induces NF-kB activation and macrophage recruitment to limit tissue invasion, assuring host survival and promoting development into the chronic bradyzoite form (50). The experimentally confirmed TRAF6-binding motifs in RON10 and GRA15 might support this process by connecting TRAF6-dependent IL-1 signalling with NF-kB activation. However, proving the functional relevance of these motifs would require further experimentation. Even though the RON6 peptide did not show binding to the TRAF6 MATH domain, it might still display binding in cells via multivalent interactions involving its 17 putative TRAF6-binding motif matches and TRAF6 trimers.

Our filtered dataset of 24,291 motif matches within 295 Toxoplasma proteins still likely contains many non-functional motif matches. More sophisticated computational approaches for match filtering would require machine learning on larger sets of validated motif instances, which, at the moment, are not available for *Toxoplasma* and other parasites. Conservation of motif matches is often used as an additional line of evidence for the prioritisation of likely functional motif matches. Unfortunately, Toxoplasma secreted proteins tend to be more novel, less conserved overall and display longer regions of disorder (Fig. S1A) (59). This makes it difficult to build good sequence alignments to compute common motif conservation metrics like in (65). Nevertheless, we included the assessment of the presence of our motif matches in different *Toxoplasma* strains and the few closely related parasite species as supplementary conservation metrics (Table S1), which could be useful when working with different strains or in understanding motif variation and evolution.

This study represents the first comprehensive survey of short linear motifs in *Toxoplasma gondii* or any other Apicomplexan parasite. There have been other efforts to predict and validate short linear motifs in pathogenic proteins (32,66). However, they focus on only a subset of proteins or specific motifs, or other pathogens such as viruses (61). Identification of novel motif instances in parasites such as Toxoplasma also enables refinement of current motif type annotations, as highlighted by some examples described in this study.

Our systematic approach for predicting instances of known motif types in Toxoplasma secreted proteins indicates that there might be a substantial number of motifs in *Toxoplasma* to be characterised, also in non-secreted proteins, e.g. motifs that participate in endogenous processes for parasite motility, cytoskeletal rearrangements, and parasite development. Thus, this dataset represents a valuable resource for hypothesis generation and design of downstream experiments to study molecular mechanisms of *Toxoplasma gondii* infection and biology. Structural models of predicted domain-motif interactions could further be used to design disruptive mutations to validated predicted modes of binding (4). Identified and experimentally validated pathogen-host motif-domain interactions can form the basis for the development of therapeutic strategies aimed at perturbing these interactions and thereby interfering with infection processes (4,10). Motif discovery in pathogens remains a much unexplored area of research, but holds great promise for improved understanding of infection mechanisms and the better design of effective treatments for parasite infection.

## Acknowledgements

We are thankful to members of the Gibson team for discussions at various stages of this work. We also express our gratitude to the curators and contributors of the ELM and the ToxoDB resources. This project was partly funded by the Deutsche Forschungsgemeinschaft (DFG, German Research Foundation) – Project-ID 449991970 granted to K.Lu.

## Author contributions

J.A.V., T.G. conceptualised the study, J.A.V. produced code and performed computational analyses, K.R., K.La., A.B. performed experiments, J.A.V. wrote the manuscript. J.A.V., T.G., K.R., K.La., K.Lu. edited the manuscript, T.G. and K.Lu. provided funding.

## Data and Code Availability

The motif discovery pipeline, processing and analysis code is made available on GitHub.(https://github.com/JesnsAV/Toxo_Motifs)

## Materials and Methods

### Retrieval of sequence, structure and motif data

*Toxoplasma* strain proteomes, as well as those of five closely related organisms from the Sarcocystidae clade, were retrieved from the ToxoDB resource (40). The fasta files of the annotated proteomes were collected for the following organisms: *Toxoplasma gondii* GT1 (Type I), *T. gondii* MAS and *T. gondii* ME49 (Type II and reference strain), *T. gondii* RUB and *T. gondii* VEG (Type III), *T. gondii* VAND, *Hammondia hammondii, Neospora caninum, Besnoitia besnoitii, Cystoisospora suis* Wien I and *Sarcosystis neurona* SN3. Using the ToxoDB search strategies, we retrieved mass spectrometry proteomic peptide mapping data, phosphosites, and cellular location predictions from the HyperLOPIT experiment (39) for the proteins of *T. gondii* ME49. All available AlphaFold database (AFDB) (67) predicted structures for *Toxoplasma gondii* ME49 were retrieved from the Google Cloud Console public-datasets-deepmind-alphafold-v4, taxonomy id: 50877 (02.05.2025). The latest dataset of motif classes was downloaded from the ELM resource (30.04.2025) (23). The file contained each motif code name, its regular expression (REGEX) and supplementary information. We did manual annotation of the motif classes to determine if their taxonomic range included vertebrates and *Toxoplasma*, i.e. if the domain was annotated in ELM to be present in the respective organism for interaction with the motif, ultimately excluding motif classes of organisms such as plants, insects and fungi. We also added a manual annotation on whether the motif is functional in an extracellular context and if it requires a phosphorylation or modification to interact with its domain partner. We retrieved available T. gondii domain mappings from UniProt (29.07.2025) (43), which came from Pfam, PROSITE and SMART.

### Motif prediction and structural assessments

We determined the presence of motif matches in the *T. gondii* ME49 by doing a baseline REGEX search. To assess the accessibility of these motif predictions, two approaches were selected: one that takes into account the intrinsic disorder of the sequence and another that evaluates the accessibility of the motif from predicted AlphaFold DB (AFDB) structures. The IUPred3 software (41) was used to calculate the disorder probability of the full proteome of *T. gondii* ME49. Once a REGEX match was determined, the IUPred3 score from all motif residues was retrieved and averaged. Complementarily, using available AFDB structures, we took the pLDDT scores from all motif residues and averaged them (68). We calculated residue accessibility using DSSP (Dictionary of Secondary Structure of Proteins) software (69) on each AFDB structure. The individual residue scores were then standardised according to (70) to obtain relative surface accessibility scores (RSA). We then took the RSA scores of all motif residues and averaged them. We determined if a motif match overlapped with known domain boundaries from UniProt by using positional information.

### Motif organism presence assessment

A reciprocal best hit search was carried out using BLAST to construct sets of orthologous sequence groups among *Toxoplasma* strains and related species. These sequence groups contained the sequences with the lowest e-value from each organism’s proteome, so alignments contained a standard number of sequences and were easily comparable. Multiple sequence alignments were produced for each sequence group using ClustalO with default parameters (71). Standard conservation measurements (65) were not possible as our alignments contained only a few species. Thus, the conservation of the motifs was approximated by carrying out a presence assessment. A motif was considered present in another organism if a match of the same class was present in its sequence within a window of 15 amino acids in the alignment around the start site of the motif match of the reference *Toxoplasma* strain ME49. And then the presence proportion of the total number of both strains and species was calculated and associated with each motif match, resulting in three different scores: the proportion of strains, species and overall organisms.

### Data enrichment and filtering

The data from the ELM REGEX motifs matches, their conservation, IUPred3, pLDDT, and accessibility scores were combined into a single table together with their respective protein information on subcellular location, annotations, domain overlap, and modifications. This was carried out using their unique ToxoDB identifiers and the motif name, match number and residue positions. Data filtering to define a set of motif matches was carried out with three different strategies. Firstly, we retained motifs from proteins with at least one peptide mapped to them from retrieved mass spectrometry data, to make sure proteins were expressed. We then made use of the ELM taxonomy range by ensuring that the taxonomic range of the motif instance, if the motif has been reported in a certain group of organisms, was valid for the *Toxoplasma* matches. Only motif classes that apply to Vertebrates and Apicomplexans were kept, mainly excluding those characterised in plants, insects or fungi. The classes valid in Vertebrates were kept if the motifs were located in secreted proteins, as they would be in contact with the vertebrate host (human, a mammal or birds) cell components. The classes only valid in extracellular contexts were kept if the motifs were located in microneme proteins, as among secreted proteins, they would be the ones in this location. We applied structural filters in two steps; only motif matches with IUPred3 scores above 0.4 were considered, following disordered region filters in (72). Secondly, for motifs in proteins with available AFDB structures, we retained the ones with a pLDDT<=80 (68) and an average RSA>= 0.25 (73). For motifs without AFDB structures, we retained only the ones outside of known domain boundaries.

### AlphaFold modelling

The TRAF6 MATH domain and the RON10 TRAF6-binding motif were selected for structural modelling. We used a local installation of AlphaFold Multimer version 2.3.219 (74). We manually defined domain boundaries to include missing secondary structure with high pLDDT values. We extended the motif sequence by including 5 residues at each sequence end. Structural model visualisations were produced using ChimeraX-1.10 (75).

The local installation of AlphaFold Multimer version 2.3.2 was run using the following parameters and following Alphafold GitHub instructions (https://github.com/deepmind/alphafold#running-alphafold):

--model_preset=multimer \

--db_preset=full_dbs \

--max_template_date=2020-05-14 \

--num_multimer_predictions_per_model=1 \

--use_gpu_relax=True \

--

bfd_database_path=/mnt/storage/alphafold/v232/bfd/bfd_metaclust_clu_complete_id 30_c90_final_seq.sorted_opt \

--

mgnify_database_path=/mnt/storage/alphafold/v232/mgnify/mgy_clusters_2022_05.f a \

--obsolete_pdbs_path=/mnt/storage/alphafold/v232/pdb_mmcif/obsolete.dat \

--

pdb_seqres_database_path=/mnt/storage/alphafold/v232/pdb_seqres/pdb_seqres.tx t \

--template_mmcif_dir=/mnt/storage/alphafold/v232/pdb_mmcif/mmcif_files \

--uniprot_database_path=/mnt/storage/alphafold/v232/uniprot/uniprot.fasta \

--uniref90_database_path=/mnt/storage/alphafold/v232/uniref90/uniref90.fasta \

--uniref30_database_path=/mnt/storage/alphafold/v232/uniref30/UniRef30_2021_03

\

--use_precomputed_msas=True

### Protein Expression and Purification

A gene encoding the human TRAF6 C-domain (residues 346-504, UniProt ID Q9Y4K3) was synthesised (GeneArt, Thermo Fisher Scientific) and subcloned into the pCoofy4 plasmid (76) containing an N-terminal His6-MBP-3C fusion tag. For the human TRAF6 C-domain (InterPro: IPR008974) gene, the native DNA sequence was used. The resulting bacterial expression construct was freshly transformed into *E. coli* BL21(DE3) codon + RIL cells (Stratagene). Precultures were grown overnight at 37°C in LB medium supplemented with 30 µg/ml Kanamycin and 35 µg/ml Chloramphenicol and used to inoculate the large-scale expression cultures. 10 ml preculture was added to 1 litre of TB-FB medium supplemented with 30 µg/ml Kanamycin and 35 µg/ml Chloramphenicol. The cultures were grown at 37°C until the OD600 reached ∼0.8, after which the temperature was reduced to 18°C and expression of the hTRAF6 C-domain was induced with 0.5 mM IPTG. After overnight expression at 18°C, the cultures were harvested by centrifugation (30 min, 5000 g, 4°C), and the pellets were flash-frozen in liquid nitrogen and stored at -80°C until the start of the protein purification.

The cells were lysed by 5 passages through a microfluidiser device, followed by centrifugation (30 min, 140.000 g, 4°C). The cleared lysate was loaded onto a 5 ml Protino Ni-NTA column (Macherey-Nagel) pre-equilibrated with 50 mM Tris-HCl, pH 8.0, 500 mM NaCl and 20 mM imidazole. After loading, the Ni-NTA column was washed with equilibration buffer until the A280 nm signal returned to baseline and eluted in a 60 ml gradient going from 0 to 100% of Ni-NTA elution buffer (50 mM Tris-HCl, pH 8.0, 500 mM NaCl, 350 mM imidazole). The Ni-NTA elution fractions were analysed by SDS-PAGE, and the fractions containing the His6-MBP-3C-hTRAF6 C-domain fusion protein were pooled. 2 mg of His6-tagged HRV 3C protease was added, and the sample was dialysed overnight at 4°C to IEX buffer (25 mM HEPES, pH 7.5, 100 mM NaCl, 20 mM imidazole). The next day, the dialysed sample was loaded onto a 5 ml Protino Ni-NTA column (Macherey-Nagel) coupled to a 5 ml HiTrap Q HP anion exchange column (Cytiva), both equilibrated with IEX buffer. The untagged hTRAF6 C-domain protein was present in the flow-through of the coupled columns, which was concentrated to 5 ml and injected into a HiLoad 16/600 Superdex 75 pg size exclusion chromatography column (Cytiva) pre-equilibrated with 25 mM HEPES pH 7.5, 150 mM NaCl and 1 mM DTT. The elution fractions containing pure hTRAF6 C-domain protein were pooled and concentrated to 2.1 mg/ml (111 µM). The final hTRAF6 C-domain protein samples were aliquoted, flash-frozen in liquid nitrogen and stored at -80°C until further usage. The identity of the hTRAF6 C-domain protein was verified by mass spectrometry (EMBL Proteomics Core Facility), and the oligomerisation state (monomeric) and absence of aggregates were confirmed by size exclusion chromatography column coupled to multi-angle light scattering (SEC-MALS). The stability of the hTRAF6 C-domain protein in the storage buffer was assessed by nano-Differential Scanning Fluorimetry (nano-DSF; Tm ∼ 46.8 °C).

### Binding assays

We selected 4 motifs from 4 different *Toxoplasma* proteins with different AF model scores for carrying out binding assays as described in Results, together with the human MAVS TRAF6 motif, being selected as a control (56). Peptides of 15 amino acids containing the motifs were defined, as well as versions containing E>S point mutations at motif position +3. The peptides were labelled with 5-Carboxyfluorescein (5-Fluo) at the C-terminus by Biosyntan GmbH (https://www.biosyntan.de) to carry out Microscale thermophoresis (MST). The lyophilised peptides were resuspended in 25mM HEPES, pH 7.5 and 150mM NaCl to a concentration of 2mM, and the pH was adjusted to 7.5 when necessary. MST measurements were performed using a Monolith NT.115 (NanoTemper Technology). Two-fold serial dilutions of the hTRAF6 protein (99 μM) were mixed in a 1:1 ratio with 100nM peptides. Titration measurements were taken for all *Toxoplasma* peptides, and the human MAVS control, with two-fold serial dilutions of the hTRAF6 protein (99 μM) mixed in a 9:1 ratio with 500nM. All measurements were performed in triplicate and carried out at 25°C with an LED excitation power of 20% and a medium MST power. Finally, the data were analysed assuming a 1:1 binding model and graphed using the MO.Affinity Software (NanoTemper Technology).

## Supplementary information

**Table S1 –** Motif predictions in microneme, rhoptry and dense granule proteins

**Table S2 –** Selected TRAF6 motif-containing peptides and binding assay results

**Data S1** – MST binding extended results for the selected TRAF6 motif-containing peptides

**Supplementary Figure 1.**
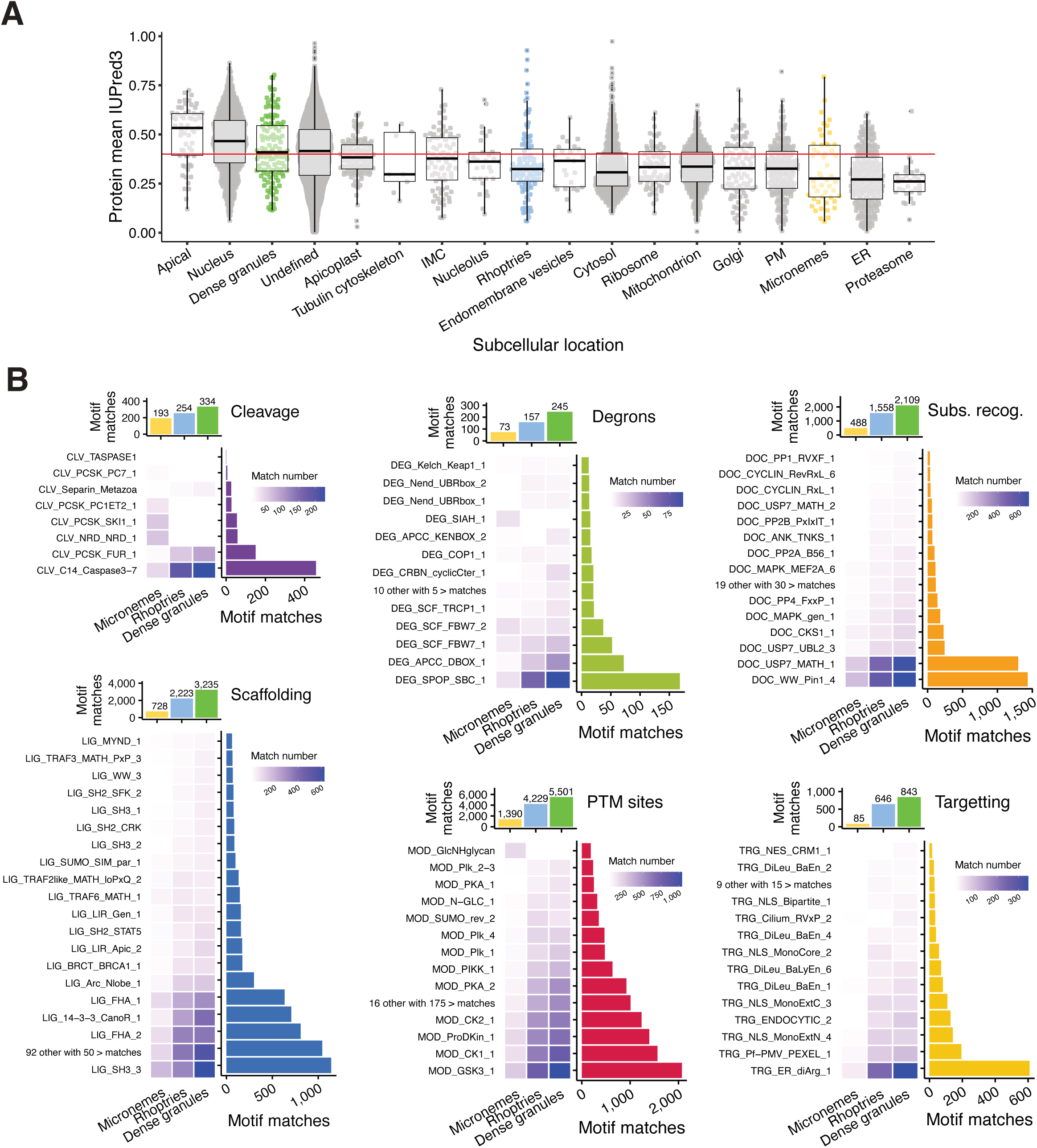
Protein disorder in different locations and matches in secreted proteins. **A**. IUPred3 mean score for proteins of the different LOPIT subcellular locations. **B**. Motif matches of the most frequent motif classes by motif.

