## Supplementary Data 1 for "Short linear motifs - Underexplored players driving *Toxoplasma gondii* infection"

### Supplementary Data - MST results

TgRON10 (peptide ID: 10784)
Sequence: 5-Fluo-TNGTPE**E**YEET-Amid

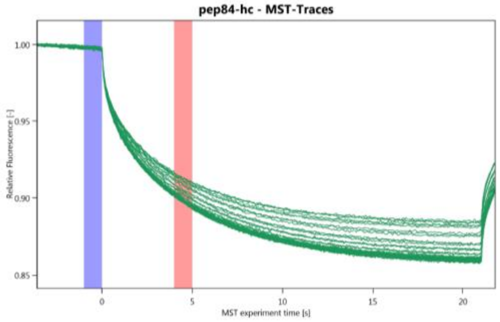


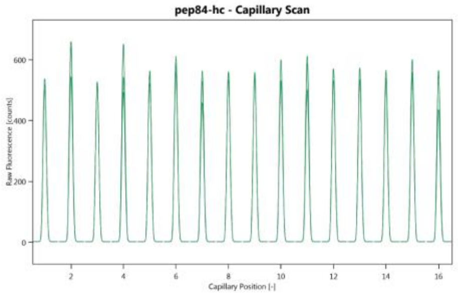


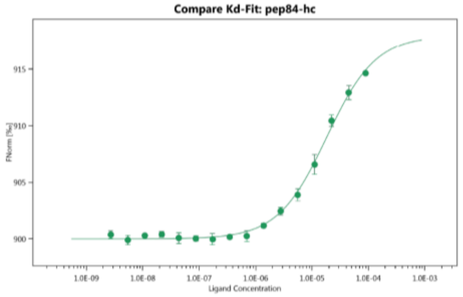


Dataset Overview

| | |
|---|---|
| Name: | hTRAF6 Tray |
| Graph Color: | • |
| Target Name: | pep84 |
| Target Concentration: | 50 nM |
| Ligand Name: | hTRAF6 |
| Ligand Concentration: | 8.91E+04 nM to 2.72 nM |
| n: | 3 |
| Comments: | |
| Excitation Power: | 20% |
| MST Power: | 40% |
| Temperature: | 25.0°C |
| Kd: | 1.8214E-05 |
| Kd Confidence: | ± 1.4653E-06 |
| Response Amplitude: | 17.985219 |
| TargetConc: | 5E-08[Fixed] |
| Unbound: | 899.98 |
| Bound: | 917.97 |
| Std. Error of Regression: | 0.3041618 |
| Reduced $\chi^2$: | 2.0670054 |
| Signal to Noise: | 63.516276 |

**Conclusion: binds (Kd = 18.2μM)**

TgRON10 mutant (peptide ID: 10785)
Sequence: 5-Fluo-TNGTPE**S**YEET-Amid

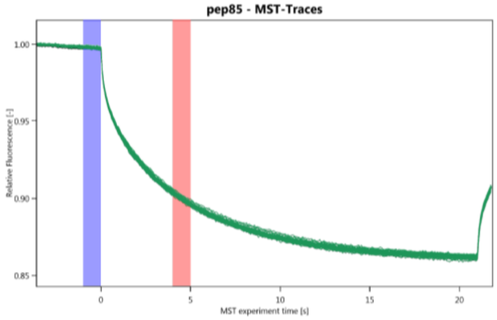


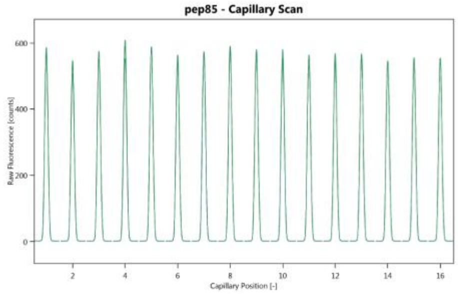


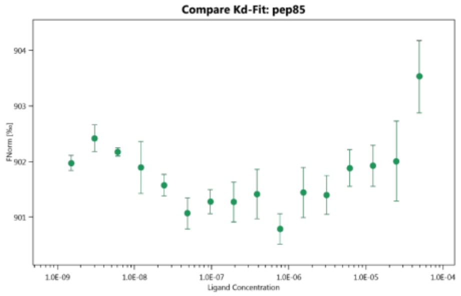


Dataset Overview

| | |
|---|---|
| Name: | hTRAF6 Tray |
| Graph Color: | • |
| Target Name: | pep85 |
| Target Concentration: | 50 nM |
| Ligand Name: | hTRAF6 |
| Ligand Concentration: | 49.5 μM to 0.00151 μM |
| n: | 3 |
| Comments: | |
| Excitation Power: | 20% |
| MST Power: | 40% |
| Temperature: | 25.0°C |
| Kd: | |
| Kd Confidence: | |
| Response Amplitude: | |
| TargetConc: | 5E-08[Fixed] |
| Unbound: | |
| Bound: | |
| Std. Error of Regression: | |
| Reduced $\chi^2$: | |
| Signal to Noise: | |

Conclusion: no binding

### TgRON6 (peptide ID: 10786)

Sequence: 5-Fluo-HERAPG**E**GGHE-Amid

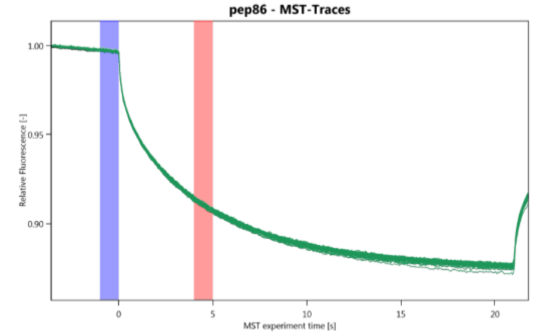


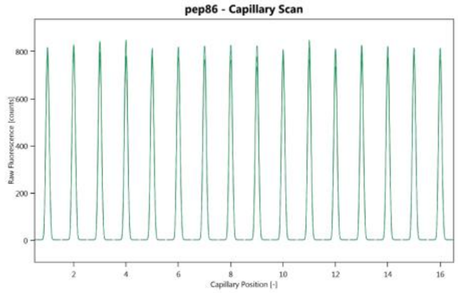


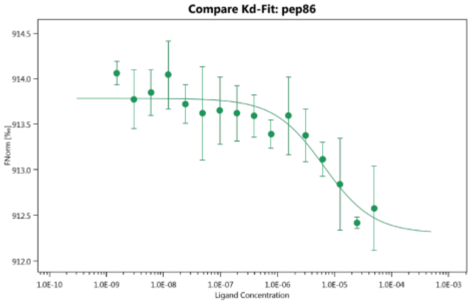


Dataset Overview

| | |
|---|---|
| Name: | hTRAF6 Tray |
| Graph Color: | • |
| Target Name: | pep86 |
| Target Concentration: | 50 nM |
| Ligand Name: | hTRAF6 |
| Ligand Concentration: | 49.5 µM to 0.00151 µM |
| n: | 3 |
| Comments: | |
| Excitation Power: | 20% |
| MST Power: | 40% |
| Temperature: | 25.0°C |
| Kd: | 6.4052E-06 |
| Kd Confidence: | ± 2.854E-06 |
| Response Amplitude: | 1.479834 |
| TargetConc: | 5E-08[Fixed] |
| Unbound: | 913.78 |
| Bound: | 912.3 |
| Std. Error of Regression: | 0.16039139 |
| Reduced $\chi^2$: | 1.3669731 |
| Signal to Noise: | 9.9107359 |

Conclusion: no binding

### TgRON6 mutant (peptide ID: 10787)

Sequence: 5-Fluo-HERAPG**S**GGHE-Amid

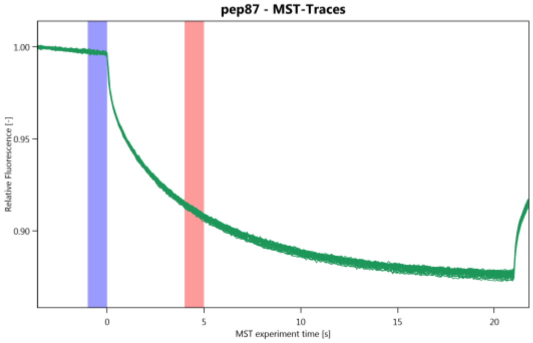


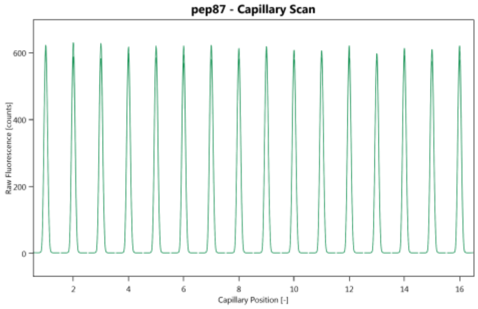


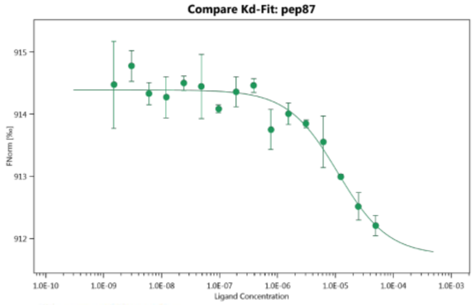


Dataset Overview

| | |
|---|---|
| Name: | hTRAF6 Tray |
| Graph Color: | • |
| Target Name: | pep87 |
| Target Concentration: | 50 nM |
| Ligand Name: | hTRAF6 |
| Ligand Concentration: | 49.5 µM to 0.00151 µM |
| n: | 3 |
| Comments: | |
| Excitation Power: | 20% |
| MST Power: | 40% |
| Temperature: | 25.0°C |
| Kd: | 1.0955E-05 |
| Kd Confidence: | ± 4.1052E-06 |
| Response Amplitude: | 2.6642171 |
| TargetConc: | 5E-08[Fixed] |
| Unbound: | 914.39 |
| Bound: | 911.72 |
| Std. Error of Regression: | 0.20315397 |
| Reduced $\chi^2$: | 2.1338393 |
| Signal to Noise: | 14.086992 |

Conclusion: no binding

TgGRA7 (peptide ID: 20443)
Sequence: 5-Fluo-LETEPD**E**QEEV-Amid

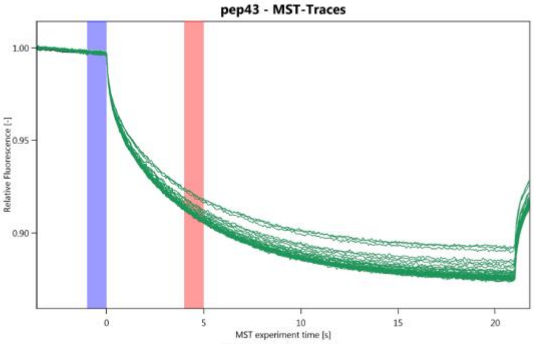


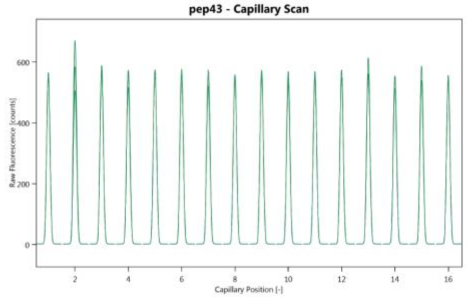


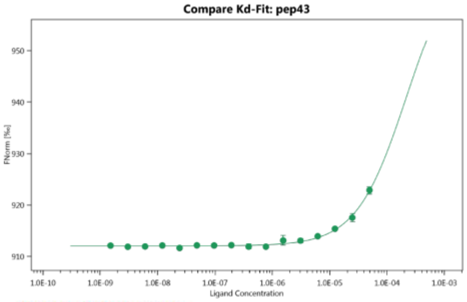


Dataset Overview

| | |
|---|---|
| Name: | hTRAF6 Tray |
| Graph Color: | • |
| Target Name: | pep43 |
| Target Concentration: | 50 nM |
| Ligand Name: | hTRAF6 |
| Ligand Concentration: | 49.5 μM to 0.00151 μM |
| n: | 3 |
| Comments: | |
| Excitation Power: | 20% |
| MST Power: | 40% |
| Temperature: | 25.0°C |
| Kd: | 0.00021695 |
| Kd Confidence: | ± 0.0001174 |
| Response Amplitude: | 57.732424 |
| TargetConc: | 5E-08[Fixed] |
| Unbound: | 912.05 |
| Bound: | 969.78 |
| Std. Error of Regression: | 0.30432054 |
| Reduced χ²: | 0.6919831 |
| Signal to Noise: | 203.78043 |

Conclusion: very weak binding

TgGRA7 mutant (peptide ID: 20444)
Sequence: 5-Fluo-LETEPD**S**QEEV-Amid

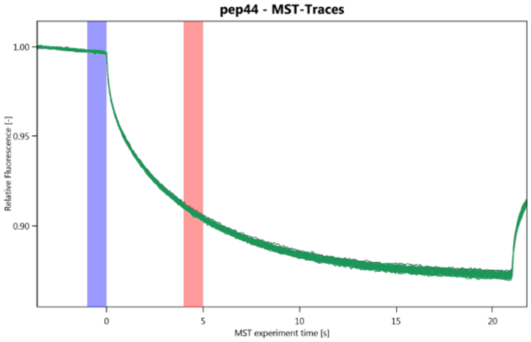


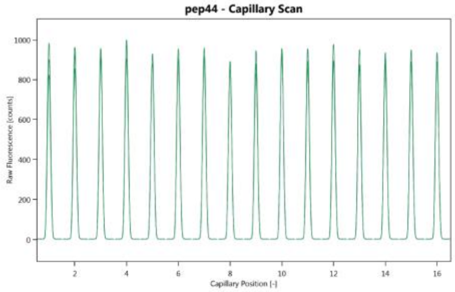


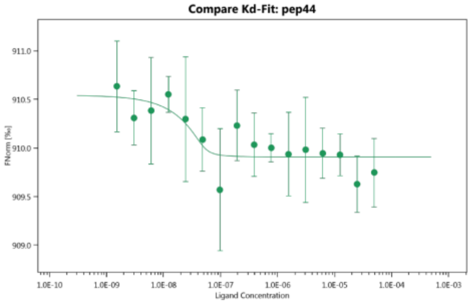


Dataset Overview

| | |
|---|---|
| Name: | hTRAF6 Tray |
| Graph Color: | • |
| Target Name: | pep44 |
| Target Concentration: | 50 nM |
| Ligand Name: | hTRAF6 |
| Ligand Concentration: | 49.5 μM to 0.00151 μM |
| n: | 3 |
| Comments: | |
| Excitation Power: | 20% |
| MST Power: | 40% |
| Temperature: | 25.0°C |
| Kd: | 1.6596E-09 |
| Kd Confidence: | ± 6.4862E-09 |
| Response Amplitude: | 0.63489257 |
| TargetConc: | 5E-08[Fixed] |
| Unbound: | 910.54 |
| Bound: | 909.91 |
| Std. Error of Regression: | 0.18643298 |
| Reduced χ²: | 0.31914315 |
| Signal to Noise: | 3.6580654 |

Conclusion: no binding

TgGRA15 (peptide ID: 20445)
Sequence: 5-Fluo-QPQVPG**E**NSYY-Amid

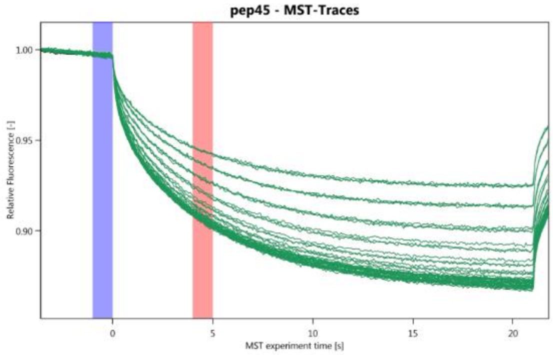


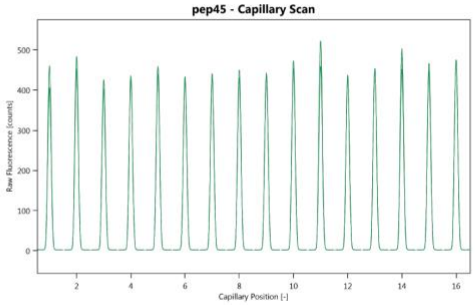


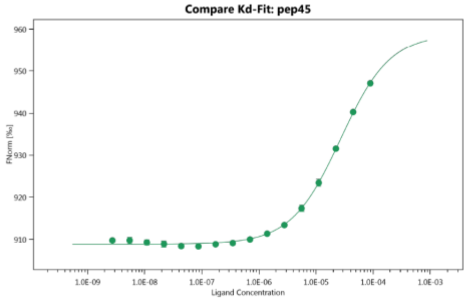


Dataset Overview

| | |
|---|---|
| Name: | hTRAF6 Tray |
| Graph Color: | • |
| Target Name: | pep45 |
| Target Concentration: | 50 nM |
| Ligand Name: | hTRAF6 |
| Ligand Concentration: | 8.91E+04 nM to 2.72 nM |
| n: | 3 |
| Comments: | |
| Excitation Power: | 20% |
| MST Power: | 40% |
| Temperature: | 25.0°C |
| Kd: | 2.6913E-05 |
| Kd Confidence: | ± 1.4303E-06 |
| Response Amplitude: | 49.948887 |
| TargetConc: | 5E-08[Fixed] |
| Unbound: | 908.86 |
| Bound: | 958.81 |
| Std. Error of Regression: | 0.47059985 |
| Reduced $\chi^2$: | 1.1021555 |
| Signal to Noise: | 114.01132 |

**Conclusion: binds (Kd = 26.9μM)**

TgGRA15 mutant (peptide ID: 20446)
Sequence: 5-Fluo-QPQVPG**S**NSYY-Amid

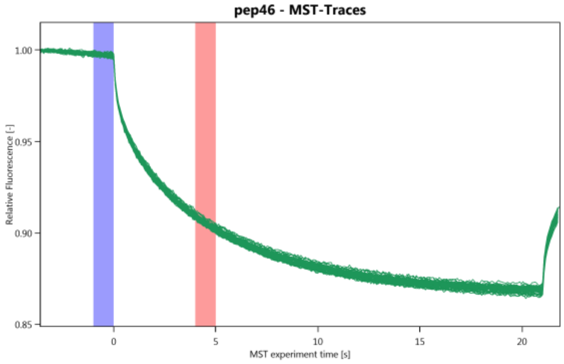


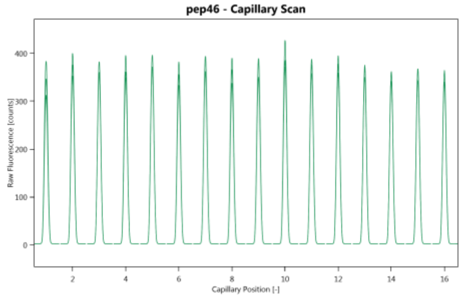


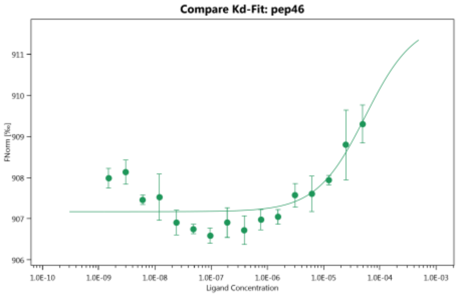


Dataset Overview

| | |
|---|---|
| Name: | hTRAF6 Tray |
| Graph Color: | • |
| Target Name: | pep46 |
| Target Concentration: | 50 nM |
| Ligand Name: | hTRAF6 |
| Ligand Concentration: | 49.5 μM to 0.00151 μM |
| n: | 3 |
| Comments: | |
| Excitation Power: | 20% |
| MST Power: | 40% |
| Temperature: | 25.0°C |
| Kd: | 5.5147E-05 |
| Kd Confidence: | ± 8.6559E-05 |
| Response Amplitude: | 4.6468734 |
| TargetConc: | 5E-08[Fixed] |
| Unbound: | 907.17 |
| Bound: | 911.82 |
| Std. Error of Regression: | 0.47406161 |
| Reduced $\chi^2$: | 4.606408 |
| Signal to Noise: | 10.529312 |

Conclusion: no binding

Human MAVS (peptide ID: 10788)
Sequence: 5-Fluo-PSHGPE**E**NEYK-Amid

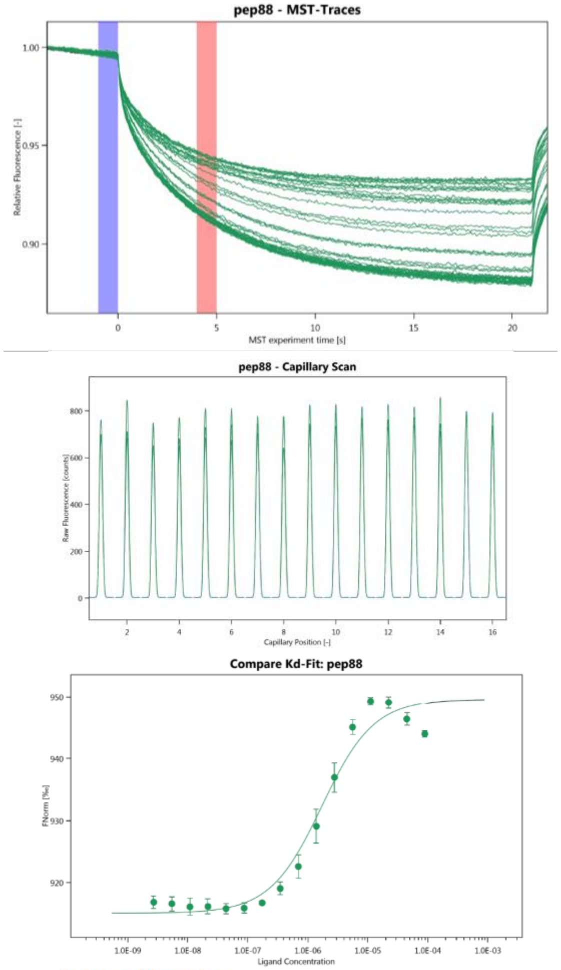


Dataset Overview

| | |
|---|---|
| Name: | hTRAF6 Tray |
| Graph Color: | • |
| Target Name: | pep88 |
| Target Concentration: | 50 nM |
| Ligand Name: | hTRAF6 |
| Ligand Concentration: | 8.91E+04 nM to 2.72 nM |
| n: | 3 |
| Comments: | |
| Excitation Power: | 20% |
| MST Power: | 40% |
| Temperature: | 25.0°C |
| Kd: | 1.6913E-06 |
| Kd Confidence: | ± 3.8719E-07 |
| Response Amplitude: | 34.532069 |
| TargetConc: | 5E-08[Fixed] |
| Unbound: | 915.06 |
| Bound: | 949.59 |
| Std. Error of Regression: | 2.5299539 |
| Reduced $\chi^2$: | 12.937886 |
| Signal to Noise: | 14.661687 |

**Conclusion: binds (Kd = 1.69μM)**

Human MAVS mutant (peptide ID: 10789)
Sequence: 5-Fluo-PSHGPE**S**NEYK-Amid

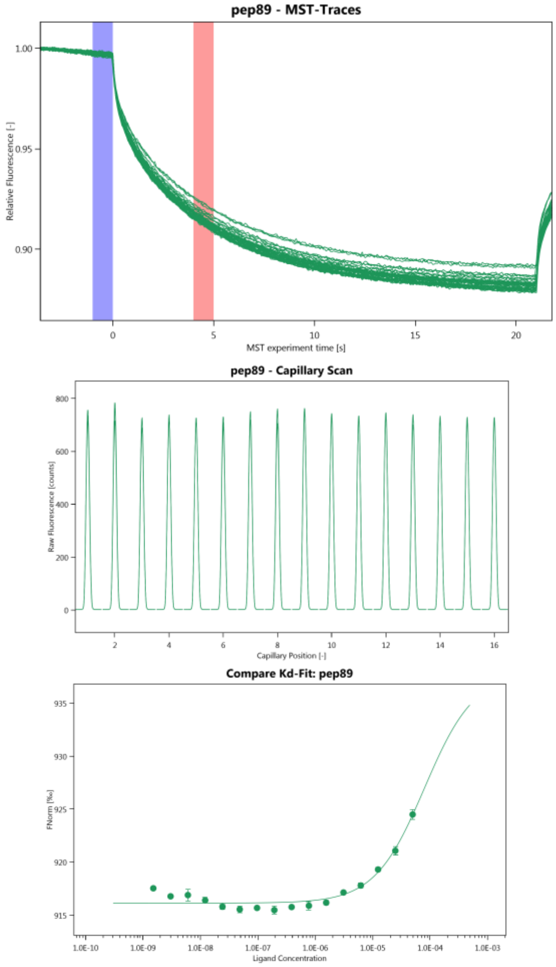


Dataset Overview

| | |
|---|---|
| Name: | hTRAF6 Tray |
| Graph Color: | • |
| Target Name: | pep89 |
| Target Concentration: | 50 nM |
| Ligand Name: | hTRAF6 |
| Ligand Concentration: | 49.5 μM to 0.00151 μM |
| n: | 3 |
| Comments: | |
| Excitation Power: | 20% |
| MST Power: | 40% |
| Temperature: | 25.0°C |
| Kd: | 7.9795E-05 |
| Kd Confidence: | ± 5.3616E-05 |
| Response Amplitude: | 21.742605 |
| TargetConc: | 5E-08[Fixed] |
| Unbound: | 916.14 |
| Bound: | 937.88 |
| Std. Error of Regression: | 0.61703124 |
| Reduced $\chi^2$: | 16.350615 |
| Signal to Noise: | 37.851089 |

Conclusion: weak binding
